# DARG: a Database of Alzheimer Related Genes in Model Organisms

**DOI:** 10.1101/2025.01.29.635533

**Authors:** Julia Y. Song, Eric Zhou, Yanhui Hu

## Abstract

Alzheimer disease is affecting a significant portion of the aging population worldwide. Despite extensive research over the years, a comprehensive understanding of its underlying mechanisms and effective treatments remains elusive. Meanwhile, animal models, such as mouse, zebrafish, Drosophila, and C. elegans, have proven invaluable in studying human diseases. In response, we have developed DARG, a Database of Alzheimer disease-related genes in model organisms, designed to bridge the gap between human geneticists investigating the molecular mechanisms of the disease and the model organisms that can be used to explore the functions of disease-associated genes. DARG allows users to search and browse Alzheimer-related human genes from various resources and datasets, identify orthologs in model organisms, and access data on their gene expression and related phenotypes in *Drosophila*.

## Introduction

The research of Alzheimer’s disease in animal models has significantly advanced our understanding of its mechanisms. The genetic manipulation to express human amyloid precursor protein (APP) and tau in various animal models, such as mouse, *zebrafish, Drosophila*, or *C. elegans*, causes key Alzheimer-related features, including age-dependent amyloid-beta aggregation and neurodegeneration, making them powerful systems for studying disease pathogenesis (Alexander *et al*. 2014; Frost *et al*. 2015; Esquerda-Canals *et al*. 2017; Thawkar and Kaur 2025). For example, transgenic mouse models, such as the APP/PS1, 3xTg-AD, and tauopathies, are widely used to study key pathological features of Alzheimer’s disease, including amyloid-beta plaque formation, tau tangles, neuroinflammation, and cognitive decline (Jankowsky et al., 2004; Oddo et al., 2003). On the other hand, *Drosophila* and *C. elegans* models remain as valuable tools in Alzheimer’s research with different advantages, including a simple nervous system, well-characterized genetics, and the ability to study aging and neurodegeneration in a short lifespan, despite their simplicity compared to vertebrate models. These models have also been used to explore genetic and environmental factors that contribute to Alzheimer’s disease. For example, studies have shown that the manipulation of various genes, including those involved in insulin signaling and autophagy, can modulate amyloid-beta toxicity and improve cognitive function in *Drosophila* model (Huang *et al*. 2019; Oyarce-Pezoa *et al*. 2023).

In addition to studying disease mechanisms, animal models are also valuable for drug screening to identify potential therapeutic compounds that reduce amyloid-beta toxicity or enhance the clearance of amyloid plaques (Van Dam and De Deyn 2011; Deshpande *et al*. 2019). For example, curcumin has been shown to alleviate Alzheimer’s disease by inhibiting inflammatory response, oxidative stress in mouse model, suggesting its potential as Alzheimer’s treatments (Zeng *et al*. 2022; Shao *et al*. 2023).

Informatics resources have been built in the past such as NIAGADS (Kuzma *et al*. 2024), AlzGene (Bertram *et al*. 2007) and AlzBase (Bai *et al*. 2016) for users to search Alzheimer related genes but most of these resources are currently not accessible anymore. To facilitate the research of Alzheimer’s disease in model organisms, we built DARG (<u>D</u>atabase of <u>A</u>lzheimer <u>R</u>elated <u>G</u>enes), an integrated resource collecting Alzheimer related human genes from various public resources, and mapped the human genes to the orthologous genes in model organisms. In addition, we also integrated the age-dependent and/or tissue specific transcriptomic data as well as phenotype annotation in nervous system from FlyBase. We believe this is a valuable resource for human geneticists investigating the disease’s molecular mechanisms as well as the researchers using the model organisms to explore the functions of disease-associated genes.

## Method

1. Retrieve human Alzheimer related genes from public resources Alzheimer related genes were collected from public resources of human disease annotation including OMIM (An Online Catalog of Human Genes and Genetic Disorders, https://omim.org/) (Amberger and Hamosh 2017), GWAS Catalog (An online catalog of human genome-wide association studies, https://www.ebi.ac.uk/gwas/) (Sollis *et al*. 2023), ClinVar (a public archive of reports of human variations classified for diseases and drug responses, with supporting evidence, https://ftp.ncbi.nlm.nih.gov/pub/clinvar/gene_condition_source_id) (Landrum *et al*. 2016), AGI (Alliance Genome Resources, https://www.alliancegenome.org/) (Alliance of Genome Resources 2024), UniProt (https://www.uniprot.org/) (UniProt 2023) and MalaCards (https://www.malacards.org/) (Rappaport *et al*. 2017) and BioLitMine (https://www.flyrnai.org/tools/biolitmine/web/) (Hu *et al*. 2020). The collected information of associated genes is processed. Both reported genes and mapped genes were extracted from GWAS catalogs. Different resources might use different gene and protein identifiers, therefore, we used our in-house ID mapping tool to map various identifiers to Entrez GeneID, and then integrated. The information about the number of publications co-citing both gene and Alzheimer’s disease is also retrieved from BioLitMine (Hu *et al*. 2020). The genes supported by multiple resources are ranked “high” while the genes from the reported gene column of GWAS, and/or the genes studies in two or more publications of Alzheimer focus are assigned “moderate” rank. All the other genes are assigned “low” rank. We filtered out pseudo genes, ncRNAs, etc. to focus on protein-coding genes.
2. Map human Alzheimer related genes to model organisms The assembled gene list was compared with orthologous relationships predicted by DIOPT (Hu *et al*. 2011), a voting system for ortholog prediction with more than twenty algorithms integrated. We only selected the predictions with high and moderate rank from DIOPT and mapped Alzheimer genes to mouse, zebrafish, *Drosophila* and *C. elegans*. The highest DIOPT score and the best orthologous genes from each of the model organism are reported at DARG site.
3. Retrieve tissue and stage specific transcriptomic data and phenotype annotation about orthologous genes in model organisms Tissue and stage specific RNA-seq datasets were obtained from FlyBase ftp site (https://ftp.flybase.net/releases/FB2024_06/precomputed_files/) (Thurmond *et al*. 2019). Only the datasets of the express levels of the head samples from 1-day, 4-day and 20-day adult flies were selected. The male and female datasets were averaged and integrated. The genes involved in abnormal neuroanatomy or abnormal neurophysiology phenotype were retrieve from FlyBase (https://flybase.org/vocabularies) and integrated.
4. Gene set enrichment analysis (GSEA) GSEA was performed using PANGEA (Hu *et al*. 2023) and the gene set annotation selected are the KEGG pathway/disease annotation (Kanehisa *et al*. 2023), HGNC gene group annotation (Bruford *et al*. 2022), SLIM terms of biological process and cellular component annotation of gene ontology (Gene Ontology *et al*. 2023). Protein complex enrichment was done using COMPLEAT (Vinayagam *et al*. 2013).
5. Database URL: https://www.flyrnai.org/tools/AlzheimerGene

## Result

We collected Alzheimer associated genes from various public resources such as OMIM (Amberger and Hamosh 2017), GWAS (Sollis *et al*. 2023) and ClinVar (Landrum *et al*. 2016), and built a database for users to mine the list with ease (figure 1A). There are 2700 protein-coding genes in total collected (supplementary table 1) and we assigned confidence based on the number of resources as well as the publication counts if the gene is linked to two or more publications of Alzheimer focus. 388 (14%) genes from multiple resources are assigned high rank while 870 (32%) are assigned moderate rank (figure 1B). Studying the function of Alzheimer genes in animal models have proven to be important to advance our understanding about the molecular mechanism of Alzheimer disease(Alexander *et al*. 2014; Frost *et al*. 2015; Esquerda-Canals *et al*. 2017; Thawkar and Kaur 2025). To facilitate such studies, we also mapped the human genes to the major model organisms using DIOPT (Hu *et al*. 2011). For example, 2670 (99%) human genes can be mapped to mouse while 2560 (95%) human genes can be mapped to Zebrafish. On the other hand, 2070 (77%) human genes are mapped to *Drosophila* while 1997 (74%) human genes are mapped to *C. elegans*. It is expected that a larger portion of genes are conserved cross species between genetically closer species, nevertheless, most of genes can be studied using any of these model organisms (figure 1C). We examined the gene expression in *Drosophila* nervous system based on transcriptome datasets available at FlyBase (Thurmond *et al*. 2019) and observed that for 2047 human genes, the corresponding *Drosophila* ortholog is also expressing in nervous system. In addition, the mutant alleles of more than 50% of these *Drosophila* orthologs for 1072 human genes also have abnormal neuroanatomy or neurophysiology phenotype, making *Drosophila* a great model to study the function of Alzheimer genes.

**Figure 1.**
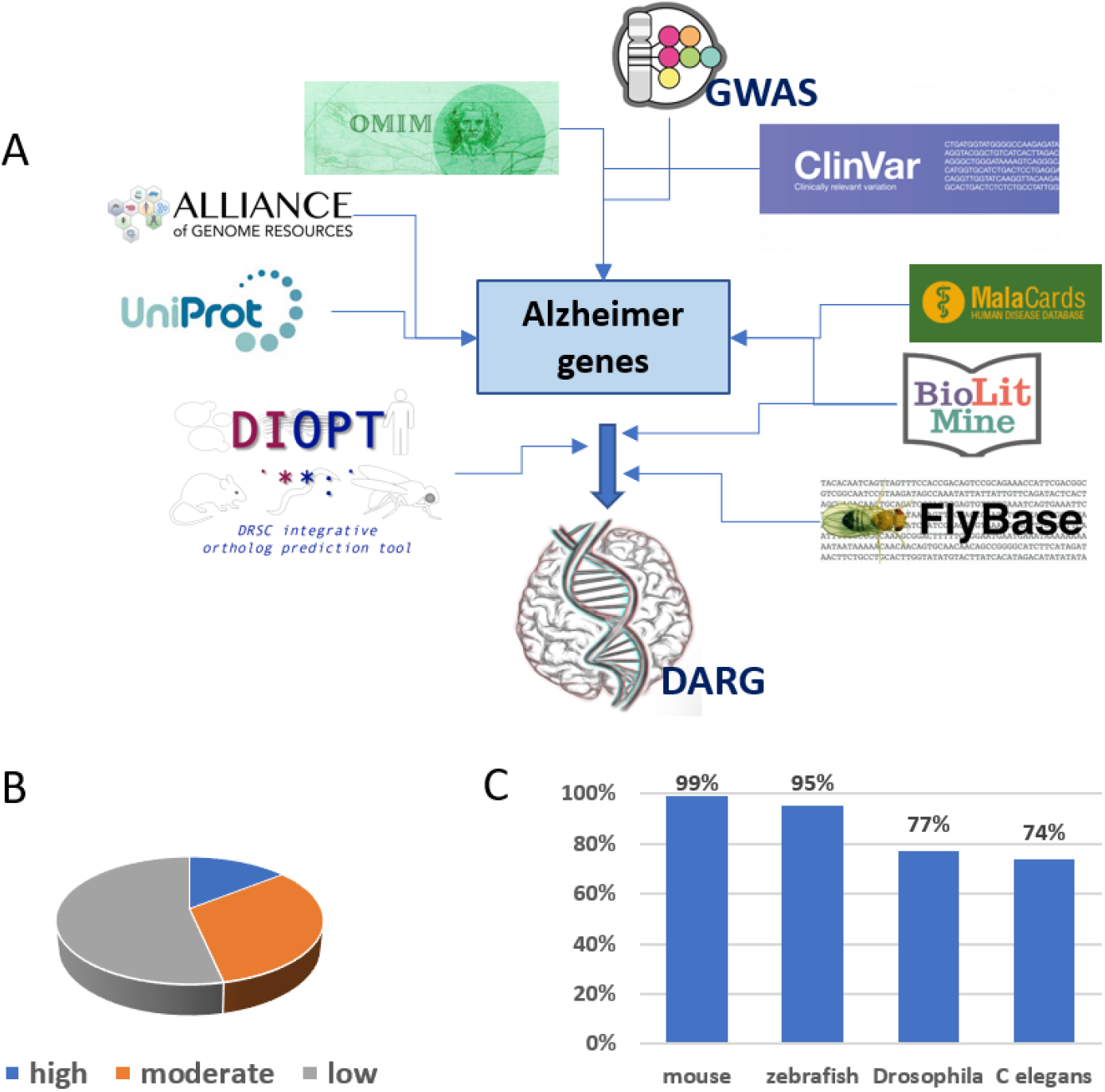
Build DARG database of Alzheimer genes. **A.)** workflow of database building **B.)** High rank is assigned to genes from multiple sources while moderate rank genes are obtained from one resource with at least 2 papers co-cited with Alzheimer or from GWAS reported gene list. **C.)** Percent of genes conserved in various model organisms.

To validate the list assembled and understand the biological context underlining this gene list, we did gene set enrichment analysis using PANGEA (Hu *et al*. 2023). The top gene sets from KEGG annotation (Kanehisa *et al*. 2023) that are highly over-represented among the high rank Alzheimer genes include “Neurotrophin signaling pathway”, “Apoptosis”, “Lipid and atherosclerosis”, “Alzheimer disease”, while the top gene sets from gene group annotation of HGNC (HUGO gene nomenclature Committee) (Bruford *et al*. 2022) enriched are “Dopamine receptors”, “Neurotrophins”, “Caspases”, “Apolipoproteins”, which are expected (figure 2). In addition, the top gene sets from the GO (gene ontology) biological process annotation include “aging”, “protein maturation”, “autophagy” while the top GO cellular component terms are “synapse”, “cell junction”, and “mitochondrion” (Gene Ontology *et al*. 2023) (figure 3). The GSEA results using the full list also show similar results with less significant p values/fold changes (data not shown), which reflects the current research in the literature that the role of high rank genes in Alzheimer are more extensively studied than moderate and low rank genes in the list. The GSEA results reflect the current understanding about Alzheimer’s disease in terms of the biological processes involved, the protein functions and the sub-cellular localizations, therefore, it validates the gene list assembled. Proteins and genes usually do not work alone, therefore, we also examined the protein complexes enriched among the high-ranking Alzheimer genes using COMPLEAT (Vinayagam *et al*. 2013) (figure 4). For example, the protein complex HC4886 is identified in this analysis, which is a protein complex of ten members that positively regulates apoptosis while HC8780, a protein complex of five members, regulates neuronal synaptic plasticity. The complex analysis results demonstrate the more detailed molecular mechanisms that underline the related biological processes for Alzheimer’s disease.

**Figure 2.**
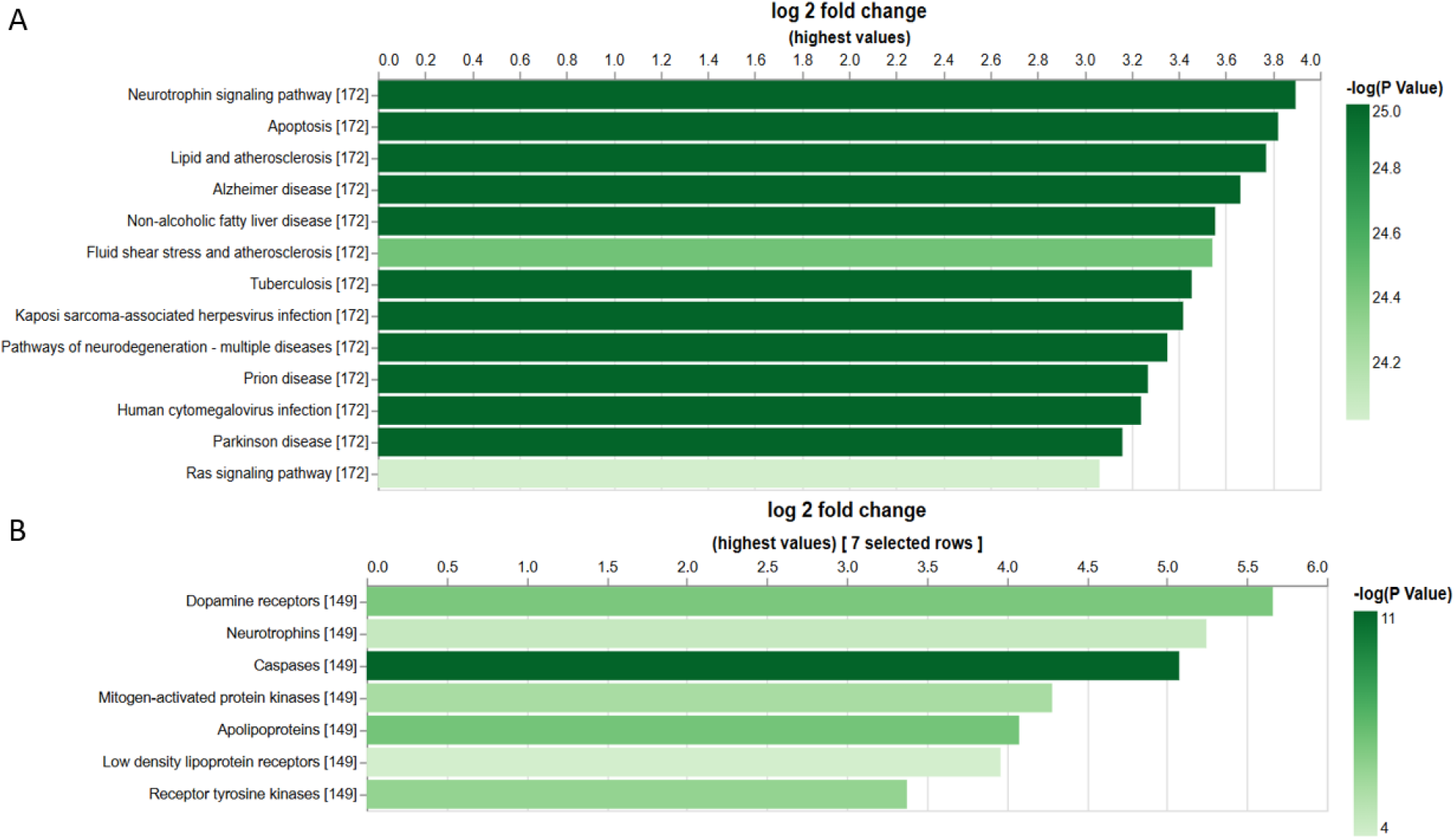
Gene set enrichment analysis using PANGEA. Gene set used are from **A.)** KEGG annotation **B.)** HGNC gene group annotation. The bar height reflects fold enrichment while the darkness reflects enrichment P value.

**Figure 3.**
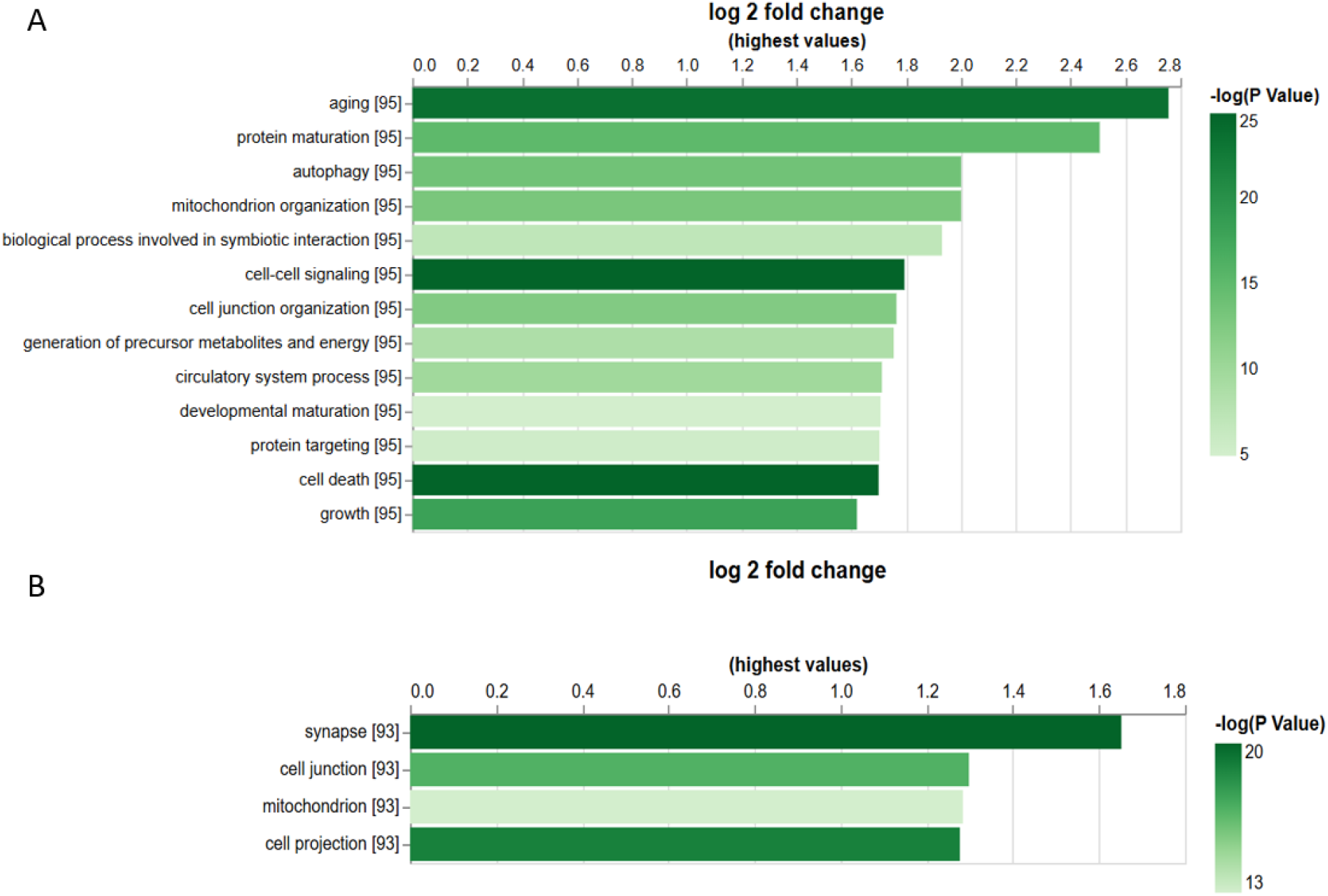
Gene set enrichment analysis using PANGEA. Gene set used are from **A.)** Biological process annotation of GO SLIM1 **B.)** Cellular component annotation of GO SLIM2. The bar height reflects fold enrichment while the darkness reflects enrichment P value.

**Figure 4.**
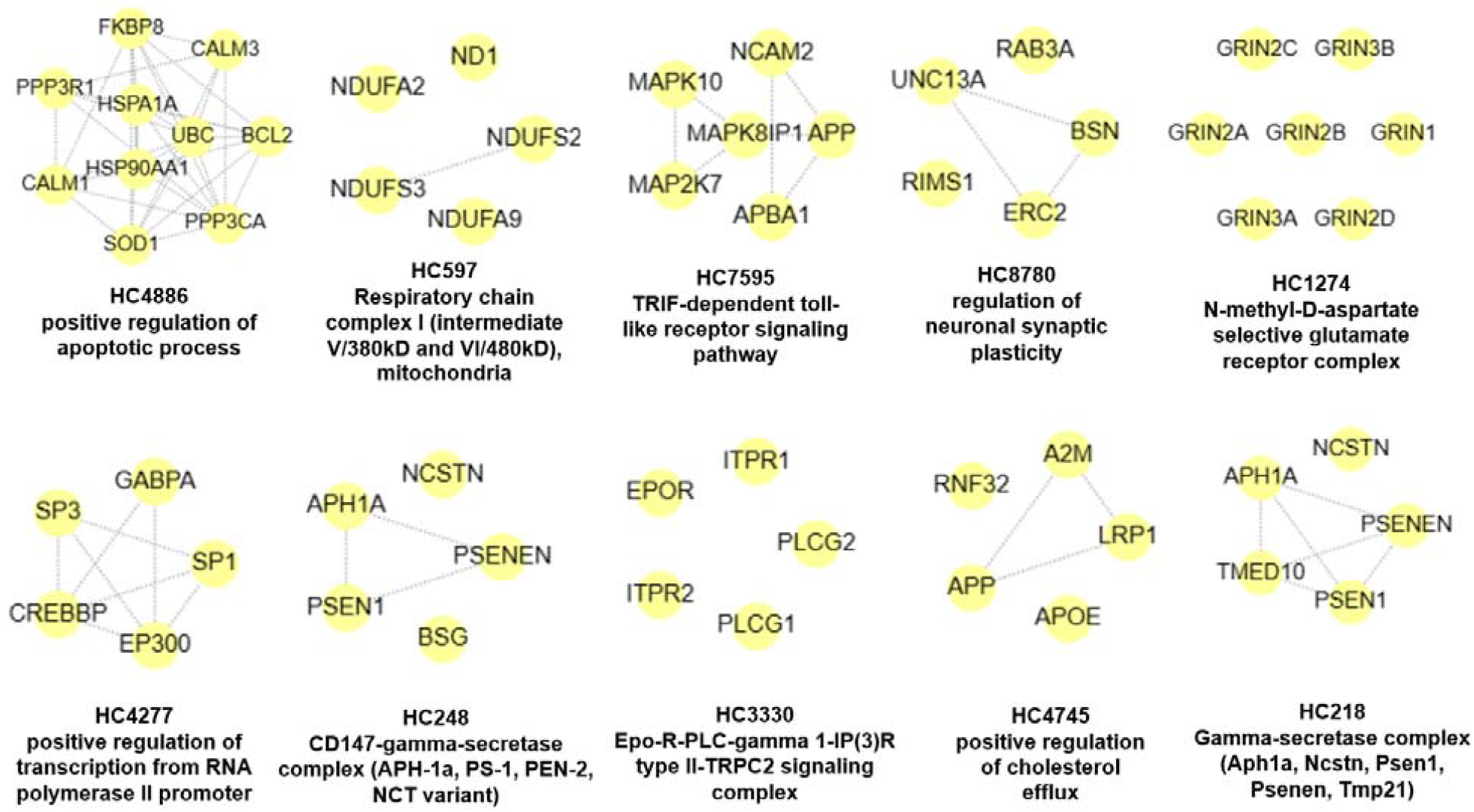
Protein complex enrichment analysis using COMPLEAT: examples of over-represented protein complexes among high-ranking Alzheimer genes.

## Discussion

The genes related to Alzheimer’s disease from different research projects are scattered in many public resources and there is an unmet need to integrate the candidate genes into a single comprehensive database. On the other hand, research on Alzheimer’s disease in various model organisms has provided valuable insights into the molecular mechanisms underlying the disease and has facilitated the discovery of potential therapeutic targets. With these in mind, we developed an integrated database of human Alzheimer candidate genes with the information of orthologous genes in major model organisms. In addition, the transcriptomic datasets from *Drosophila* nervous system of different stage as well as the information of abnormal neuronal phenotype annotation in *Drosophila* is also integrated. Users can easily obtain a comprehensive list of candidate genes and identify the subset of genes that can be further studied in *Drosophila* or other model organisms.

Despite their advantages, animal models cannot fully replicate the complexity of Alzheimer’s disease in humans. For example, *Drosophila* models have limitations due to their simpler nervous system and the lack of some Alzheimer’s disease features observed only in humans. However, ongoing research continues to refine these models and use them to better understand the molecular underpinnings of Alzheimer disease, its genetic basis, and to screen for potential therapeutic agents. We believe DARG will bridge the gap between human geneticists studying the molecular mechanisms of the disease and researchers working with model organisms to explore the molecular functions of disease-related genes.

## References

Alexander, A. G., V. Marfil and C. Li, 2014 Use of Caenorhabditis elegans as a model to study Alzheimer’s disease and other neurodegenerative diseases. Front Genet 5: 279.

Alliance of Genome Resources, C., 2024 Updates to the Alliance of Genome Resources central infrastructure. Genetics 227.

Amberger, J. S., and A. Hamosh, 2017 Searching Online Mendelian Inheritance in Man (OMIM): A Knowledgebase of Human Genes and Genetic Phenotypes. Curr Protoc Bioinformatics 58: 1 2 1–1 2 12.

Bai, Z., G. Han, B. Xie, J. Wang, F. Song et al., 2016 AlzBase: an Integrative Database for Gene Dysregulation in Alzheimer’s Disease. Mol Neurobiol 53: 310–319.

Bertram, L., M. B. McQueen, K. Mullin, D. Blacker and R. E. Tanzi, 2007 Systematic meta-analyses of Alzheimer disease genetic association studies: the AlzGene database. Nat Genet 39: 17–23.

Bruford, E. A., B. Braschi, L. Haim-Vilmovsky, T. E. M. Jones, R. L. Seal et al., 2022 The importance of being the HGNC. Hum Genomics 16: 58.

Deshpande, P., N. Gogia and A. Singh, 2019 Exploring the efficacy of natural products in alleviating Alzheimer’s disease. Neural Regen Res 14: 1321–1329.

Esquerda-Canals, G., L. Montoliu-Gaya, J. Guell-Bosch and S. Villegas, 2017 Mouse Models of Alzheimer’s Disease. J Alzheimers Dis 57: 1171–1183.

Frost, B., J. Gotz and M. B. Feany, 2015 Connecting the dots between tau dysfunction and neurodegeneration. Trends Cell Biol 25: 46–53.

Gene Ontology, C., S. A. Aleksander, J. Balhoff, S. Carbon, J. M. Cherry et al., 2023 The Gene Ontology knowledgebase in 2023. Genetics 224.

Hu, Y., V. Chung, A. Comjean, J. Rodiger, F. Nipun et al., 2020 BioLitMine: Advanced Mining of Biomedical and Biological Literature About Human Genes and Genes from Major Model Organisms. G3 (Bethesda) 10: 4531–4539.

Hu, Y., A. Comjean, H. Attrill, G. Antonazzo, J. Thurmond et al., 2023 PANGEA: a new gene set enrichment tool for Drosophila and common research organisms. Nucleic Acids Res 51: W419–W426.

Hu, Y., I. Flockhart, A. Vinayagam, C. Bergwitz, B. Berger et al., 2011 An integrative approach to ortholog prediction for disease-focused and other functional studies. BMC Bioinformatics 12: 357.

Huang, Y., Z. Wan, Z. Wang and B. Zhou, 2019 Insulin signaling in Drosophila melanogaster mediates Abeta toxicity. Commun Biol 2: 13.

Kanehisa, M., M. Furumichi, Y. Sato, M. Kawashima and M. Ishiguro-Watanabe, 2023 KEGG for taxonomy-based analysis of pathways and genomes. Nucleic Acids Res 51: D587–D592.

Kuzma, A., O. Valladares, E. Greenfest-Allen, H. Nicaretta, M. Kirsch et al., 2024 NIAGADS: A Comprehensive National Data Repository for Alzheimer’s Disease and Related Dementia Genetics and Genomics Research. medRxiv.

Landrum, M. J., J. M. Lee, M. Benson, G. Brown, C. Chao et al., 2016 ClinVar: public archive of interpretations of clinically relevant variants. Nucleic Acids Res 44: D862–868.

Oyarce-Pezoa, S., G. G. Rucatti, F. Munoz-Carvajal, N. Sanhueza, W. Gomez et al., 2023 The autophagy protein Def8 is altered in Alzheimer’s disease and Abeta42-expressing Drosophila brains. Sci Rep 13: 17137.

Rappaport, N., M. Twik, I. Plaschkes, R. Nudel, T. Iny Stein et al., 2017 MalaCards: an amalgamated human disease compendium with diverse clinical and genetic annotation and structured search. Nucleic Acids Res 45: D877–D887.

Shao, S., X. Ye, W. Su and Y. Wang, 2023 Curcumin alleviates Alzheimer’s disease by inhibiting inflammatory response, oxidative stress and activating the AMPK pathway. J Chem Neuroanat 134: 102363.

Sollis, E., A. Mosaku, A. Abid, A. Buniello, M. Cerezo et al., 2023 The NHGRI-EBI GWAS Catalog: knowledgebase and deposition resource. Nucleic Acids Res 51: D977–D985.

Thawkar, B., and G. Kaur, 2025 The current models unravel the molecular mechanisms underlying the intricate pathophysiology of Alzheimer’s disease using zebrafish. Methods Cell Biol 192: 17–31.

Thurmond, J., J. L. Goodman, V. B. Strelets, H. Attrill, L. S. Gramates et al., 2019 FlyBase 2.0: the next generation. Nucleic Acids Res 47: D759–D765.

UniProt, C., 2023 UniProt: the Universal Protein Knowledgebase in 2023. Nucleic Acids Res 51: D523–D531.

Van Dam, D., and P. P. De Deyn, 2011 Animal models in the drug discovery pipeline for Alzheimer’s disease. Br J Pharmacol 164: 1285–1300.

Vinayagam, A., Y. Hu, M. Kulkarni, C. Roesel, R. Sopko et al., 2013 Protein complex-based analysis framework for high-throughput data sets. Sci Signal 6: rs5.

Zeng, Y., Y. Lv, M. Hu, F. Guo and C. Zhang, 2022 Curcumin-loaded hydroxypropyl-beta-cyclodextrin inclusion complex with enhanced dissolution and oral bioavailability for epilepsy treatment. Xenobiotica 52: 718–728.

